# Functional data analysis techniques to improve the generalizability of near-infrared spectral data for monitoring mosquito populations

**DOI:** 10.1101/2020.04.28.058495

**Authors:** Pedro M. Esperança, Dari F. Da, Ben Lambert, Roch K. Dabiré, Thomas S. Churcher

## Abstract

Near infrared spectroscopy is increasingly being used as an economical method to monitor mosquito vector populations in support of disease control. Despite this rise in popularity, strong geographical variation in spectra has proven an issue for generalising predictions from one location to another. Here, we use a functional data analysis approach—which models spectra as smooth curves rather than as a discrete set of points—to develop a method that is robust to geographic heterogeneity. Specifically, we use a penalised generalised linear modelling framework which includes efficient functional representation of spectra, spectral smoothing and regularisation. To ensure better generalisation of model predictions from one training set to another, we use cross-validation procedures favouring smoother representation of spectra. To illustrate the performance of our approach, we collected spectra for field-caught specimens of *Anopheles gambiae* complex mosquitoes – the most epidemiologically important vector species on the planet – in two sites in Burkina Faso. Using these spectra, we show how models trained on data from one site can successfully classify morphologically identical sibling species in another site, over 250km away. Whilst we apply our framework to species prediction, our unified statistical framework can, alternatively, handle regression analysis (for example, to determine mosquito age) and other types of multinomial classification (for example, to determine infection status). To make our methods readily available for field entomologists, we have created an open-source R package `mlevcm`. All data used is publicly also available.

## 1 Introduction

Mosquito-borne diseases such as malaria, dengue and yellow fever are responsible for huge suffering, death and impose a considerable economic burden in Sub-Saharan Africa, Asia, and Latin America (Sachs and Malaney, 2002; WHO, 2019). The World Health Organization estimated 228 million cases of malaria alone in 2018 resulting in approximately 405,000 deaths. Malaria is transmitted from person to person by female mosquitoes of the *Anopheles* genus. Insecticides which kill mosquitoes, either incorporated into bednets or sprayed on walls, are the most effective method of controlling the disease and prevent millions of cases each year (Bhatt et al., 2015). However, differences in behaviour between mosquito species and the rise of insecticide resistance mean that control interventions increasingly need to be tailored to the local mosquito population. Factors such as species composition, the level of mosquito infection and age distribution in the mosquito population constitute an important direct measure of the efficacy of disease control interventions.

Unfortunately, there is no easy way to cheaply monitor mosquito populations in the field. Molecular techniques, like polymerase chain reaction, are required to determine mosquito species and infection status, which are laborious and require highly trained staff, well equipped laboratories and expensive reagents. By killing mosquitoes, the main effect of insecticides is to reduce mosquito lifespan, shifting the age distribution towards younger mosquitoes. Insecticide resistance may reduce the killing effect of insecticides, increasing the average age in mosquito populations supposedly controlled by insecticides. Therefore, methods to monitor the local age distribution in mosquito populations are critical for knowing whether insecticides remain effective. Yet, there are currently no fast, inexpensive methods for accurately surveying the age distribution of mosquito populations.

Near-infrared spectroscopy (NIRS) is a new, rapid, reagent-free and non-destructive scanning technique, which can determine the species of morphologically indistinguishable mosquitoes, approximate mosquito age and the presence of malaria and dengue infections (Esperança et al., 2018; Lambert et al., 2018; Mayagaya et al., 2009; Ong et al., 2020; Sikulu et al., 2010; Sikulu-Lord et al., 2016). The instrument is portable and battery powered, which means scanning can take place in remote locations. The scanning procedure itself does not require expensive reagents, specialised lab-trained staff or non-portable laboratory equipment and is extremely simple: mosquitoes are killed, placed under a light probe, and scanned to produce a spectrum within seconds. Previous work has successfully predicted characteristics of interest (age, species, infectiousness) from near-infrared (NIR) spectra, using only relatively basic machine learning models based on Partial Least Squares (PLS) regression (Gerlach et al., 1979).

Despite their success in predicting characteristics of interest within a given population of mosquitoes, it has been documented that these methods cannot predict these between populations: that is, models trained to predict (say) age in population A cannot predict age in population B (Lambert et al., 2018). This site specificity is not typically reported in NIRS studies of mosquitoes (e.g. Esperança et al., 2018) and means the reported performance of the method may exceed that in the field. Whilst the exact origins of this between-site performance are unknown, many possible factors may contribute. In a ‘typical’ NIRS study, performance of the method is evaluated by predicting mosquito characteristics in independent test sets. Whilst different mosquitoes may be used in the training and testing sets, they come from the same set of mosquitoes, potentially from the same mother, and are kept in identical rearing conditions. These shared factors mean mosquitoes comprising the test set are much more alike those in the training set than any wild-caught specimens. Part of these issues could be addressed by training models using F0 or F1 mosquitoes derived from collected individuals – although, admittedly, such restrictions on experimental practices would limit the usefulness of NIRS. There is, hence, a demand for machine learning methods that are robust to differences between laboratory training and wild testing sets.

In this article, we use a functional data analysis (FDA) framework to build NIR spectra-based machine learning models which maintain predictive accuracy between populations of mosquitoes. In FDA, individual data points are modelled as originating from (noisy) sampling of unobserved smooth, continuous functions at discrete intervals along them (Ramsay and Silverman, 2005). Since the observed NIR spectra already appear quite smooth (see Fig. 1), this suggests that an FDA approach should be applicable. The statistical framework for functional data has been developed in the past two decades and has been used for both regression (i.e. continuous response) and classification (i.e. categorical response) problems. The flexibility of FDA means that modern techniques such as efficient function representations, smoothing, penalised estimation and dimension reduction can be accommodated seamlessly—all of which we explore here (Morris, 2015; Ramsay and Silverman, 2002, 2005; Reiss et al., 2017; Wang et al., 2016).

**Figure 1:**
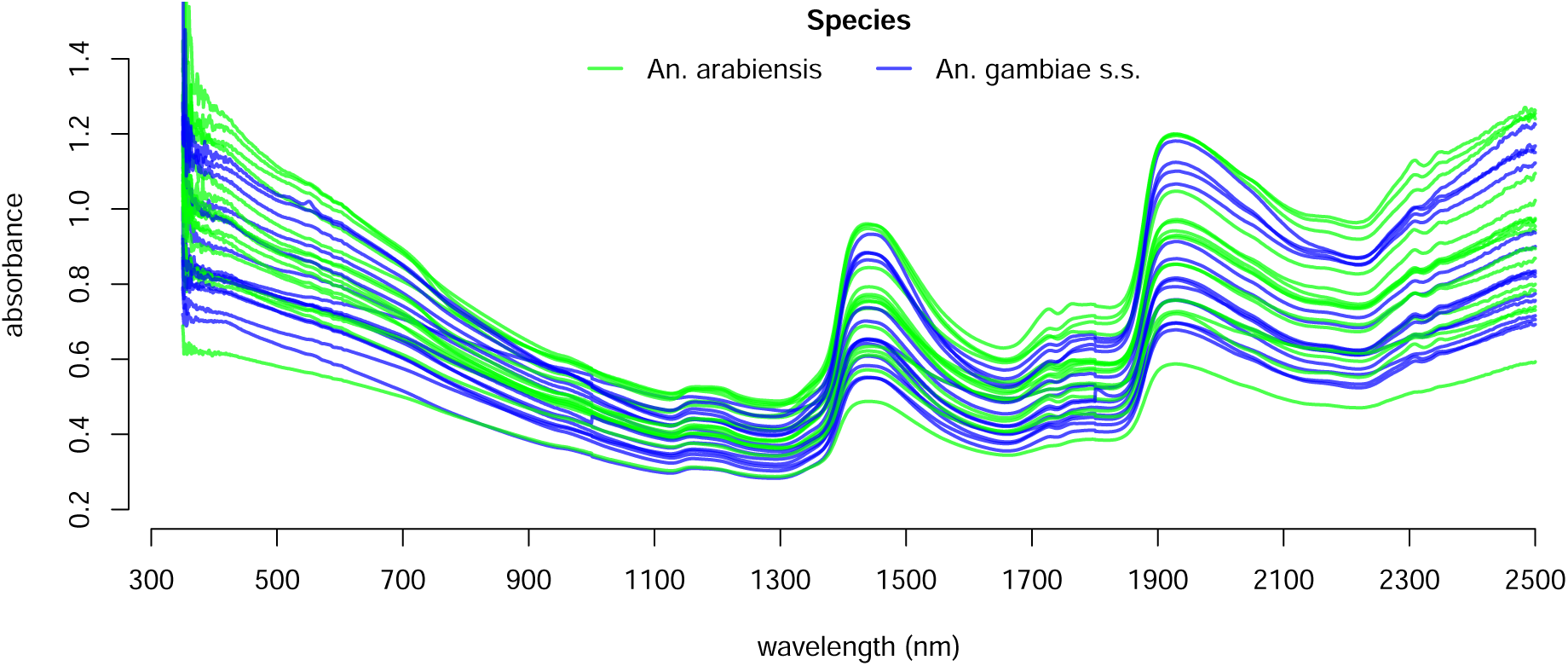
The spectra of 30 mosquitoes within the *Anopheles gambiae s.l.* (sensu lato) species complex, each sampled at discrete wavelengths in the interval [350, 2500]. Green lines show *Anopheles arabiensis* whilst blue lines show *Anopheles gambiae s.s.* (sensu stricto).

To demonstrate the utility of our approach, we generated training and testing samples that mimicked how NIRS could be used in the field: we collected wild *Anopheles gambiae s.l.* mosquito larvae from two locations in Burkina Faso, separated by 283km, which were reared then scanned using near-infrared spectrometers in the laboratory. After scanning, the mosquitoes were killed and their species was determined by PCR. (Specimens were either *An. gambiae s.s.* or *An. arabiensis*, which are morphologically identical species that have epidemiologically important differences in ecology.) We then show that our FDA-based approach trained using paired species-spectra data from each location in isolation can predict the species of individual specimens in the other. To encourage others to replicate and build on our analysis, we make all data (Esperança, 2019a) and code (Esperança, 2019b) publicly available.

## 2 Methods

### 2.1 Mosquito data

To train and test our machine learning algorithm, we collected mosquito larvae from two locations in Burkina Faso: Klesso, in the southwest of the country, near Bobo-Dioulasso; and Longo, in the Hauts-Basins region approximately 283km away. Adult mosquitoes were reared from field collected larvae (F0 generation) or from wild, naturally fed mosquitoes caught resting in the eaves of houses which were allowed to lay eggs which were then reared to adult (F1 generation). All mosquitoes were kept in similar ‘laboratory colony’ conditions and killed using chloroform four days after emergence. Mosquitoes were scanned using a LabSpec4 Standard-Res i (standard resolution, integrated light source) near-infrared spectrometer and a bifurcated reflectance probe mounted 2mm from a spectralon white reference panel (ASD Inc. (company), 2020). Absorbance was recorded across 350nm–2500nm of the electromagnetic spectrum. Specimens were laid on their side under the focus of the light probe and spectra were recorded with RS3 spectral acquisition software (ASD Inc. (company), 2020), which recorded the average spectra from 20 scans. The light probe was centred on the head and thorax region of the mosquito and each mosquito was scanned up to 4 times, picking the mosquito and replacing on their opposite side after each scan. The average number of scans per mosquito is 2.5 (75% with 2 scans, 24% with 4 scans, 1% with 1 or 3 scans). The mean absorbance across the multiple scans was then used in the analyses (Figure 1). After mosquitoes have been scanned, species was determined by Polymerase Chain Reaction (Fanello et al., 2002). This resulted in 224 spectra samples in Klesso (50 *An. arabiensis* and 174 *An. gambiae s.s.*) and 126 Longo (61 *An. arabiensis* and 65 *An. gambiae s.s.*).

### 2.2 Statistical Methods

Our approach follows a unified framework for functional data analysis (FDA; Ramsay and Silverman, 2005). As pre-processing, we represent spectra efficiently using basis functions (§2.2.1) and perform smoothing to eliminate measurement noise (§2.2.2). To classifying mosquito species we use a regularised, generalised linear model framework (§2.2.3) with dimension reduction (§2.2.4), and use a cross-validation procedure to optimise hyperparameters (§2.2.5). All variables that we use to describe our method are summarised in Table 1.

**Table 1:**
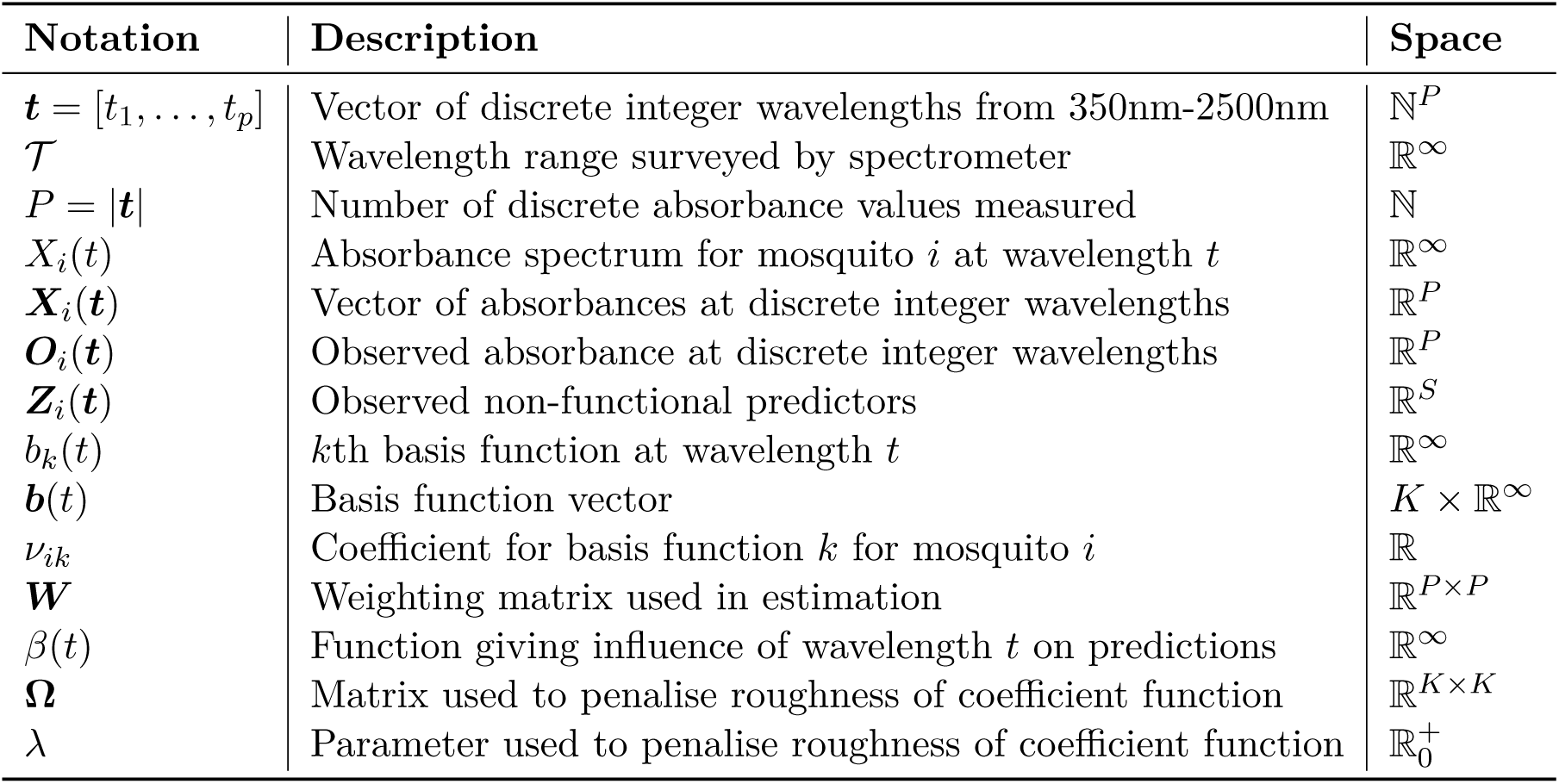
Notation. Description of variables and their ranges.

#### 2.2.1 Spectra as functional data

Mosquito spectra can be viewed as smooth curves or functions sampled at discrete wavelengths in the near-infrared (NIR) region of the electromagnetic spectrum (Figure 1). Therefore, Functional Data Analysis (FDA) can be used to represent these data (Ramsay and Silverman, 2005).

Let *X*(*t*) represent the true, underlying absorbance spectrum of a mosquito as a function of wavelength *t* ∈ 𝒯. For each spectra sample, absorbances are reported at all integer wavelengths between 350nm and 2500nm. We denote the vector of absorbances for mosquito *i* by ***X***_*i*_(***t***) = [*X*_*i*_(350), …, *X*_*i*_(2500)], *i* ∈ [1:*N*].

We follow the standard theoretical framework in FDA, which assumes that functions are real-valued and belong to a Hilbert space containing square-integrable functions over the observed range of wavelengths (Febrero-Bande et al., 2017; Reiss et al., 2017).

#### 2.2.2 Basis function representation and spectra smoothing

Basis functions provide an accurate and efficient way of representing complex functions as combinations of simpler functions, and constitute also to a natural framework for smoothing.

##### Representation

We express spectra as a linear combination of a set of *K* basis functions,

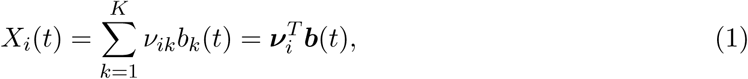

where ***b***(*t*) = [*b*_1_(*t*), …, *b*_*K*_ (*t*)]^*T*^ is a basis function vector, with *b*_*k*_(*t*) denoting the *k*th basis function evaluated at wavelength *t*; and ***ν***_*i*_ = [*ν*_*i*1_, …, *ν*_*iK*_]^*T*^ is a basis coefficient vector, with *ν*_*ik*_ denoting the *k*th basis function coefficient for the *i*th mosquito spectra. The basis coefficients are estimated from the data, as detailed below, while the basis functions can be either data-dependent (e.g. principal components) or data-independent (e.g. B-splines and wavelets).

B-splines are a natural choice of basis system for NIR spectra. These are constructed from piecewise polynomial functions, typically of low degree *n*, with continuous derivatives up to derivative degree *n* − 1, which makes them appealing both theoretically and computationally (de Boor, 2001; Eubank, 1999; Green and Silverman, 1994; Silverman, 1985). We use cubic B-splines (*n* = 3), which are sufficiently flexible to represent spectra accurately. Figure 2 illustrates how spectral data can be represented using cubic B-splines.

**Figure 2:**
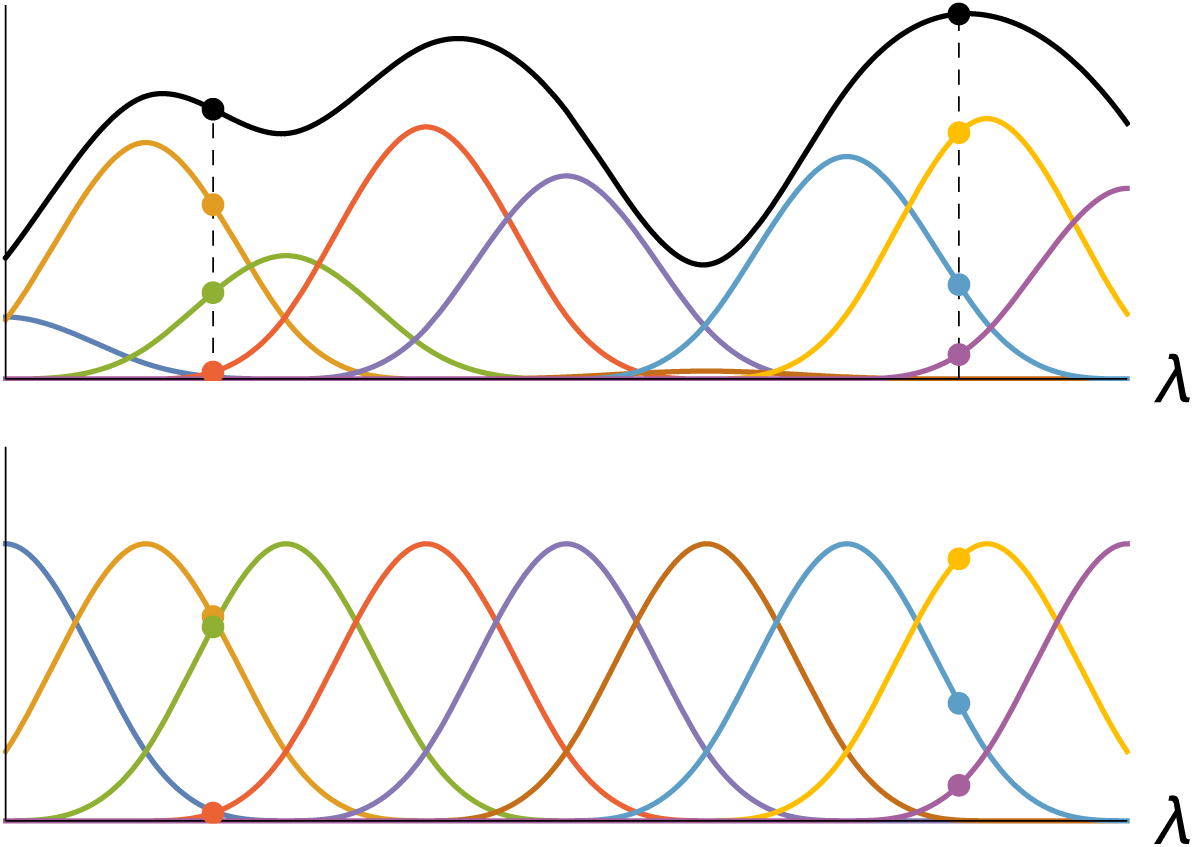
Representing near-infrared spectra using basis functions. Bottom: a number of unweighted cubic B-spline basis functions (coloured lines); the sets of points show the largest components of the basis function design matrix in two of its rows (corresponding to two measured wavelengths). Top: The absorbance spectra (black line) at two wavelengths (black points) is obtained by summing the contributions from weighted individual splines (coloured lines) as in (1).

##### Smoothing

Spectral observations are subject to measurement error due to imperfections in the spectrometer’s detection sensors (see Figure 1). Poor signal-to-noise ratios make it harder to avoid overfitting—especially, if noise varies between training and test sets. The measurement error is assumed to be Gaussian and independent for each sample *i* and wavelength *t*,

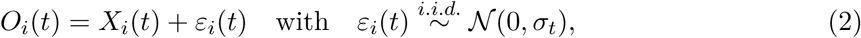

where *O*_*i*_(*t*) and *X*_*i*_(*t*) represent, respectively, the observed noisy measurements and the underlying unobserved functional process, and **Σ** = diag(*σ*_1_, …, *σ*_*P*_) represents a diagonal covariance matrix to allow for heteroscedastic measurement noise (Ramsay and Silverman, 2005).

We estimate the unobserved absorbance ***X***_*i*_(***t***) from the vector of observations ***O***_*i*_(***t***). Using the basis representation of *X*_*i*_(*t*) in (1), the measurement error model in (2) reduces to a linear regression model, 𝔼[***O***_*i*_(***t***)] = ***Bν***, where ***B*** = [***b***(*t*_1_), …, ***b***(*t*_*P*_)]^*T*^ is the *P* × *K* basis function design matrix and ***ν*** is the basis coefficient vector which can be estimated via least squares.

To correct for heteroscedastic measurement noise we introduce a *P* ×*P* weight matrix ***W*** = **Σ**^−1^. Additionally, we penalise discontinuous jumps in consecutive basis coefficient values through a *K* × *K* regularisation matrix **Ω**, and estimate *ν* by solving

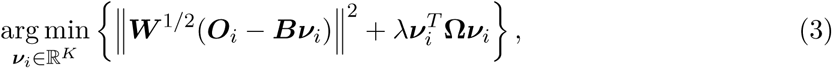

where || · || denotes the *L*_2_ norm. The *K* × *K* penalty matrix **Ω** has elements (*k, l*) equal to 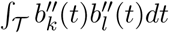, such that 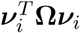 approximates the curvature of *X*_*i*_(*t*) as measured by the integrated squared second derivative, 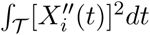 (Cardot et al., 2003; Eubank, 1999; Green and Silverman, 1994; Marx and Eilers, 1999; Ramsay and Silverman, 2005). The criterion (3) therefore enforces smoothness by penalising roughness in the least-squares estimate of *X*_*i*_(*t*). The penalty parameter *λ* ≥ 0 regulates the degree of smoothness and is optimally chosen using cross-validation (Wahba, 1990).

#### 2.2.3 Statistical Model

##### Model specification

The exponential family of statistical models can be extended to the case of functional data, providing a comprehensible and unified framework to tackle regression and classification tasks (Cardot and Sarda, 2005; Goldsmith et al., 2011; James, 2002; Müller and Stadtmüller, 2005). These models can be written as follows:

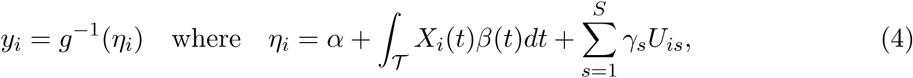

where *α* is a constant intercept; *X*_*i*_(*t*) is the spectrum of mosquito *i* (represented as a function); *β*(*t*) is a functional slope coefficient giving the influence of different wavelength regions on the response; *y*_*i*_ is a scalar response with distribution belonging to the exponential family; *U*_*is*_ is a real-valued non-functional predictor; and *γ*_*s*_ is the corresponding slope coefficient.

The invertible link function *g* relates the subject-specific mean response *µ*_*i*_ to the linear predictor *η*_*i*_ as follows: *g*(E[*y*_*i*_|*X*_*i*_(***t***)]) = *η*_*i*_, or, equivalently, *µ*_*i*_ = E[*y*_*i*_|*X*_*i*_(***t***)] = *g*^−1^(*η*_*i*_). The functional form of *g* depends on the distribution of the response, and determines the type of statistical model, as follows:

I. regression: when the response is real-valued and assumed to follow a Gaussian distribution, the link function is just the identity, that is *g*(*µ*_*i*_) = *µ*_*i*_, and so *µ*_*i*_ = *η*_*i*_, leading to the functional linear model (“*functional LM* “). This is the case when determining mosquito age.
II. classification: when the response is binary and assumed to follow a Bernoulli distribution, the link function is equal to *g*(*µ*_*i*_) = log(*µ*_*i*_*/*(1 − *µ*_*i*_)), and so 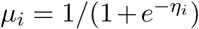, leading to the logistic-link functional generalised linear model (“*functional GLM* “). This is the case when determining mosquito species or infection.

In some applications, we may also be interested in the multi-class classification problem, for instance when information on the severity of infection is available (e.g. low, medium and high levels) or differentiating between more than two mosquito species. This corresponds to a multinomial distribution for the response, which can be reduced to a set of binary functional logistic models and therefore tackled within this same framework (McCullagh and Nelder, 1989).

##### Basis functions for *β*(*t*)

The coefficient function *β*(*t*) is modelled as a smooth function (like the spectra themselves) using cubic B-splines,

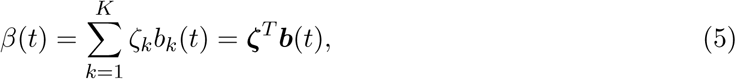

where ***b***(*t*) is the basis function vector defined as in (1); and ***ζ*** = [*ζ*_1_, …, *ζ*_*K*_]^*T*^ is a basis coefficient vector. The functional term in (4) then becomes:

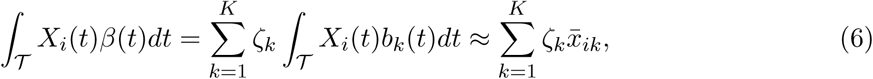

where 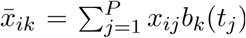 is a discretised approximation to ∫_𝒯_ *X*_*i*_(*t*)*b*_*k*_(*t*)*dt*. In this way, the functional model can be reduced to a multivariate model, for which estimation and inference procedures are well known. Notice that despite this discretisation, the functional representation of *β*(*t*) is easily recovered from (5), given estimates 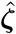of ***ζ***.

##### Model estimation

We assume independent and identically distributed pairs of observations {(*X*_*i*_(***t***), *y*_*i*_)}_*i*∈{1:*N*}_. Let ***y*** = [*y*_1_, …, *y*_*N*_]^*T*^ denote the responses vector of length *N*; and let ***X*** = [***x***_1_, …, ***x***_*N*_]^*T*^ denote the functional predictor design matrix of size *N* × *P*, where ***x***_*i*_ = [*x*_*i*1_, …, *x*_*iP*_] and *x*_*ij*_ = *X*_*i*_(*t*_*j*_) for all *i* ∈ {1:*N*} and *j* ∈ {1:*P*}. The design matrix ***X*** can denote the raw observations of the functional predictor or the corresponding smoothed version as in §2.2.2. Also, let ***B*** = [***b***(*t*_1_), …, ***b***(*t*_*P*_)]^*T*^ denote a *P* × *K* basis function design matrix, with ***b***(*t*_*j*_) = [*b*_1_(*t*_*j*_), …, *b*_*K*_ (*t*_*j*_)] for all *j* ∈ {1:*P*}. Finally, let ***Z*** = [***z***_1_, …, ***z***_*N*_]^*T*^ denote the non-functional predictor design matrix of size *N* ×*S*, where ***z***_*i*_ = [*z*_*i*1_, …, *z*_*iS*_] for all *i* ∈ {1:*N*}.

###### Estimation

Projecting the coefficient function onto the space spanned by the *K* B-splines, as defined in (5), leads to a model with design matrix ***XB*** instead of ***X***. Here we consider penalised likelihood estimation. In the linear case, the least squares solution gives the maximum likelihood estimator:

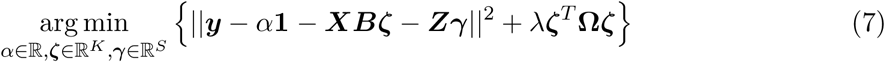

where *λ* and **Ω** are as in (3) and the term ***ζ***^*T*^ **Ω*ζ*** gives the curvature of the projected coefficient function 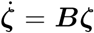, which approximates the curvature of the original coefficient function *β*(*t*) as measured by the integrated squared second derivative, ∫_𝒯_ [*β*″(*t*)]^2^*dt* (Cardot et al., 2003; Marx and Eilers, 1999; Reiss and Ogden, 2007). In the generalised linear case, the squared norm term in (7) is replaced with the negative of the model likelihood and the resulting penalised likelihood criterion is optimised (Gertheiss et al., 2013).

The problem (7) will typically be ill-posed as a result of the high dimensional nature of spectral data and the small sample sizes usually available (*N ≪ P*). Variable selection and/or dimension reduction techniques provide a solution which we explore below.

#### 2.2.4 Dimension reduction and feature selection

The projection of *β*(*t*) onto a B-spline system defined in (5) can provide some dimension reduction when *K* < *P*. However, this comes at a cost of loss of information which can lead to poor predictive performance. Here we assume a rich basis system, capable of representing spectra without any considerable loss of information. In practice, this means that the design matrix ***XB*** may still present an ill-posed problem (*K* > *N*) and further dimension reduction is then required.

We consider projecting the coefficient function onto ***D*** after the projection onto ***B***. The dimension reduction projection matrix ***D***, with dimension *Q* < *K*, is derived from the data such that ***XBD*** captures the essential features of ***XB*** (and therefore of ***X***), but in a lower dimensional space. Below we give details on two methods to compute ***D***, namely functional principal component analysis and functional partial least squares. For further comparisons of the two methods see for instance Febrero-Bande et al. (2017); Frank and Friedman (1993); Reiss and Ogden (2007).

These two projection-based dimension reduction approaches can be viewed as plug-in methods since ***D*** is independent of the parameters in the statistical model (details are given in Appendix A). Parameter estimation follows essentially the same approach as in (7), except that we now use the reduced spectral data ***XBD*** instead of ***XB***:

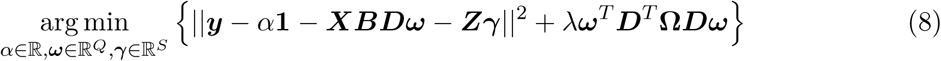

where *λ* and **Ω** are as in (3) and the term ***ω***^*T*^ ***D***^*T*^ **Ω*Dω*** gives the curvature of projected coefficient function 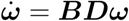. Provided that *Q* + *S* < *N*, the problem is well-posed and a solution can be found via either penalised least squares or penalised likelihood estimation as before, depending on the distribution of the response.

Importantly, note that from the estimation procedure in (8) it is possible to recover the coefficient function *β*(*t*) by first computing 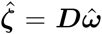 and then plugging this into (5). This tells us which regions of the spectra—in the original *P* -dimensional space—are more important.

#### 2.2.5 Cross-validation

We use a two-stage cross-validation procedure which explicitly enforces a smooth coefficient function *β*(*t*). Performance is measured by RMSD for the functional LM and by AUC for the functional GLM.

##### Datasets

To show the full potential of the techniques proposed—functional representation, smoothing, penalisation—we use two datasets. The first, called the *cross-validation dataset*, is split into training, validating and testing subsets, respectively used to train the models, cross-validate model parameters as detailed below, and estimate the generalisation error. The second, called the *alternative dataset*, is composed of a testing set which is used to evaluate the quality of predictions on slightly different samples using the model trained with the cross-validation dataset. For the application presented in this paper, this means samples collected from different regions (see §3 for more details).

##### Choosing *K*

In the first stage of cross-validation, the number of basis functions *K* used to represent the spectra is chosen to maximise the performance in a model without penalisation. To do this, *K* is decreased from *P* until the loss in accuracy exceeds the threshold *τ*_*K*_. This guarantees that there is virtually no information loss while at the same time giving the most efficient representation of the spectra. The technique provides non only an efficient representation but also a small degree of smoothing. We use *τ*_*K*_ = 0.01 in the following.

##### Choosing *Q* and *λ*

In the second stage of cross-validation, the number of PCA/PLS components *Q* and the penalty parameter *λ* are chosen jointly to give the smoothest coefficient function whose predictive performance is within a margin *τ*_*λ,Q*_ of the predictive performance of the optimal non-penalised model, which we denote by *a*_⋆_. That is, we compute models with different combinations of parameters (*λ, Q*) and from those with acceptable performance we select the one having the smoothest coefficient function *β*(*t*), as measured by the integrated squared second derivative, *R*_*β*_ = ∫_*T*_ [*β*″(*t*)]^2^*dt*, where larger values correspond to rougher coefficient functions. Acceptable performance is defined here as an RMSD between [*a*_⋆_, *a*_⋆_ + *τ*_*λ,Q*_] in the case of the functional LM, or an AUC between [*a*_⋆_ − *τ*_*λ,Q*_, *a*_⋆_] in the case of the functional GLM. We use *τ*_*λ,Q*_ = 0.01 in the following.

##### Ensemble models

We also test the performance of ensemble models where the error rate is averaged over a set of models, chosen as follows: first, we select the top *n*_a_ models that perform within a margin *τ*_*λ,Q*_ of the optimal non-penalised model (similarly to the procedure used to choose *λ* and *Q*); and from this set of acceptable models we select the smoothest *n*_e_ models. We use *n*_a_ = 25 and *n*_e_ = 5 in the following.

##### Cross-validation details

We average the cross-validation results over 100 randomisation of the data subsets to reduce the effect of sampling error. The proportions of observations used in each subset of the cross-validation dataset were: 50% for training, 25% for validating, and 25% for testing. We use as the benchmark the Generalised Linear Model (`GLM` in Table 2).

**Table 2:** Performance of different `GLM` models and different feature selection methods (details in §2.2) for determining mosquito species (*An. arabiensis* vs. *An. gambiae s.s.*). Measures given are: number of basis functions as a fraction of the number of predictors/wavelengths (*K/P*), number of features (*Q*), roughness of the coefficient function (*R*_*β*_), penalty parameter (*λ*), area under the ROC curve (*AUC*; 0–100), and cross-validation/alternative testing set errors (*ERR*_cv_/*ERR*_alt_, % misclassification rate with standard deviation in parenthesis). We show two sets of models: best-performing (highest *AUC*) and smoothest (lowest *R*_*β*_) as determined by their performance on the cross-validation dataset.

| PLS |  |  |  |  |  |  |  |  |  |  |  |  |  |
| --- | --- | --- | --- | --- | --- | --- | --- | --- | --- | --- | --- | --- | --- |
| | $K/P$ | best-performing | | | | | | smoothest | | | | | |
| | | $Q$ | $R_\beta$ | $\lambda$ | $AUC$ | $ERR_{cv}$ | $ERR_{alt}$ | $Q$ | $R_\beta$ | $\lambda$ | $AUC$ | $ERR_{cv}$ | $ERR_{alt}$ |
| GLM | 1 | 2 | 11.62 | — | 72 | 34 (.08) | 32 (.04) | — | — | — | — | — | — |
| sGLM | 1 | 8 | 0.04 | — | 75 | 30 (.07) | 30 (.05) | — | — | — | — | — | — |
| pGLM | 1 | 37 | 0.20 | 8.9 | 65 | 53 (.07) | 62 (.25) | 32 | 0.07 | 6.8 | 75 | 44 (.11) | 38 (.24) |
| spGLM | 1 | 19 | 0.001 | 61.1 | 76 | 30 (.08) | 24 (.05) | 20 | 0.001 | 94.4 | 76 | 30 (.08) | 25 (.05) |
| fGLM | 0.1 | 8 | 2.00 | — | 75 | 29 (.07) | 30 (.06) | — | — | — | — | — | — |
| fsGLM | 0.1 | 8 | 2.11 | — | 75 | 30 (.07) | 30 (.06) | — | — | — | — | — | — |
| fpGLM | 0.1 | 22 | 0.09 | 58.3 | 75 | 32 (.08) | 24 (.06) | 23 | 0.09 | 58.3 | 75 | 32 (.09) | 24 (.06) |
| fspGLM | 0.1 | 23 | 0.08 | 102 | 75 | 32 (.09) | 24 (.05) | 23 | 0.07 | 134 | 75 | 32 (.08) | 24 (.07) |
| PCA |  |  |  |  |  |  |  |  |  |  |  |  |  |
| | $K/P$ | best-performing | | | | | | smoothest | | | | | |
| | | $Q$ | $R_\beta$ | $\lambda$ | $AUC$ | $ERR_{cv}$ | $ERR_{alt}$ | $Q$ | $R_\beta$ | $\lambda$ | $AUC$ | $ERR_{cv}$ | $ERR_{alt}$ |
| GLM | 1 | 2 | 35.07 | — | 71 | 36 (.08) | 30 (.05) | — | — | — | — | — | — |
| sGLM | 1 | 10 | 0.18 | — | 75 | 30 (.08) | 29 (.06) | — | — | — | — | — | — |
| pGLM | 1 | 48 | 0.02 | 3.9 | 73 | 34 (.08) | 29 (.04) | 63 | 0.02 | 3.2 | 74 | 33 (.08) | 27 (.04) |
| spGLM | 1 | 31 | 0.001 | 53.4 | 76 | 29 (.08) | 26 (.04) | 27 | 0.001 | 53.4 | 77 | 29 (.08) | 26 (.04) |
| fGLM | 0.1 | 10 | 3.30 | — | 75 | 30 (.08) | 30 (.05) | — | — | — | — | — | — |
| fsGLM | 0.1 | 10 | 1.38 | — | 75 | 30 (.08) | 30 (.06) | — | — | — | — | — | — |
| fpGLM | 0.1 | 30 | 0.09 | 7.8 | 75 | 32 (.09) | 25 (.04) | 22 | 0.10 | 9.3 | 75 | 32 (.08) | 26 (.04) |
| fspGLM | 0.1 | 30 | 0.09 | 7.8 | 75 | 32 (.08) | 24 (.04) | 25 | 0.10 | 6.2 | 75 | 31 (.08) | 25 (.04) |

## 3 Results

We compare the performance of 16 different models, arising from the use of the two different dimension reduction methods (PLS and PCA) and the use, or not, of the FDA techniques presented. We will show results for a classification task, thus all models will be generalised linear models (`GLM`), prefixed as follows: `f` (e.g., `fGLM`) when making use of the functional representation in (1); `s` (e.g., `sGLM`) when making use of spectra smoothing as in (3); `p` (e.g., `pGLM`) when making use of penalisation for the coefficient function estimation as in (8). Additionally we evaluate ensembles of the smoothest models that use penalisation.

### 3.1 Improving generalisation

The techniques used—spectra smoothing, functional representation and penalised estimation of the coefficient function—all improve the AUC and test error on the testing subset of the cross-validation dataset, if only slightly, with the exception of penalisation-only (`pGLM`) with PLS reduction which introduces considerable variation in the estimates as shown by the standard error of the error rate (Table 2). More importantly, however, substantial improvement of the error rate can be observed on alternative testing set, from 32/30% for the benchmark model (`GLM`) to 24/25% for the model using all three techniques (`fspGLM`) with PLS/PCA reduction.

The `fpGLM` and `fspGLM` are the best performing models with very similar error rates on the alternative test set, showing that smoothing becomes only marginally important when a functional representation is used. This is not surprising seeing that a functional representation provides smoothing alongside dimension reduction. It is worth noting that a functional representation provides marginally better results then smoothing when only one of the techniques is used in conjunction with penalisation, which can be seen by comparing `spGLM` and `fpGLM`.

The relationship between smoothness and performance is as expected. Specifically, the smoothest models—here defined by a low value of *R*_*β*_, the roughness of the coefficient function—tend to perform better on the alternative testing set than rougher models.

The ensemble approach does not improve results w.r.t. the corresponding smoothest models with the exception of the `pGLM` with PLS reduction, where both error rate and standard deviation are improved substantially (Table 3). However, this still does not constitute an improvement w.r.t. the smoothest `GLM` (the benchmark). Additionally, the ensemble `pGLM` does not outperform `sGLM`, `fGLM` or `fsGLM`, suggesting that smoothing spectra is essential to avoid overfitting.

**Table 3:** Performance of different penalised ensemble models. See Table 2 for a description of the measures reported.

| | $K/P$ | PLS | | | | | | | PCA | | | | | | |
| --- | --- | --- | --- | --- | --- | --- | --- | --- | --- | --- | --- | --- | --- | --- | --- |
| | | $Q$ | $R_\beta$ | $\lambda$ | $AUC$ | $ERR_{cv}$ | $ERR_{alt}$ | | $Q$ | $R_\beta$ | $\lambda$ | $AUC$ | $ERR_{cv}$ | $ERR_{alt}$ | |
| pGLM | 1 | 31 | 0.09 | 4.5 | 75 | 41 (.11) | 33 (.18) |  | 60 | 0.02 | 3.3 | 74 | 33 (.08) | 27 (.04) |  |
| spGLM | 1 | 20 | 0.001 | 87.8 | 76 | 30 (.08) | 25 (.05) |  | 28 | 0.001 | 53.4 | 77 | 29 (.08) | 26 (.04) |  |
| fpGLM | 0.1 | 22 | 0.09 | 51.7 | 75 | 32 (.08) | 24 (.06) |  | 24 | 0.11 | 6.5 | 75 | 31 (.08) | 26 (.04) |  |
| fspGLM | 0.1 | 23 | 0.08 | 105 | 75 | 32 (.09) | 24 (.05) |  | 23 | 0.10 | 8.1 | 75 | 31 (.08) | 25 (.04) |  |

The optimal (i.e. lossless) functional representation affords a 90% compression level, which reduces the computational costs of computing the dimension reduction matrix ***D***. The resulting optimal number of PLS components *Q* increases although only marginally, for instance from 20 in the `spGLM` to 23 in the `fspGLM`. For the model using PCA components this number decreases from 27 to 25.

Importantly, the standard deviation of error rate on the alternative test set is left virtually unchanged or only minimally increased as a result of the three techniques used, again with the exception of the `pGLM` with PLS reduction.

In general, PLS gives slightly smaller error rates than PCA on the alternative testing set and requires a smaller number of components, thus performing better in both accuracy and efficiency.

The AUC is a less important performance measure since it is computed with the testing subset of the cross-validation dataset, while we are primarily interested in the performance of the model on the alternative testing set. Nonetheless, we also see an improvement between 3–4 p.p. in the AUC with the use of functional techniques.

### 3.2 Visualisation and Diagnostics

#### Cross-validation illustrated

The results of (*λ, Q*) cross-validation evaluated on the testing subset of the cross-validation dataset are illustrated in Figure 3. Accuracy is measured by the AUC and the models with acceptable performance, as defined in §2.2.5, are shown within the boxed area (Figure 3a). From the set of acceptable models, the smoothest or an ensemble are picked to produce the predictions on the alternative dataset, reported in Table 2.

**Figure 3:**
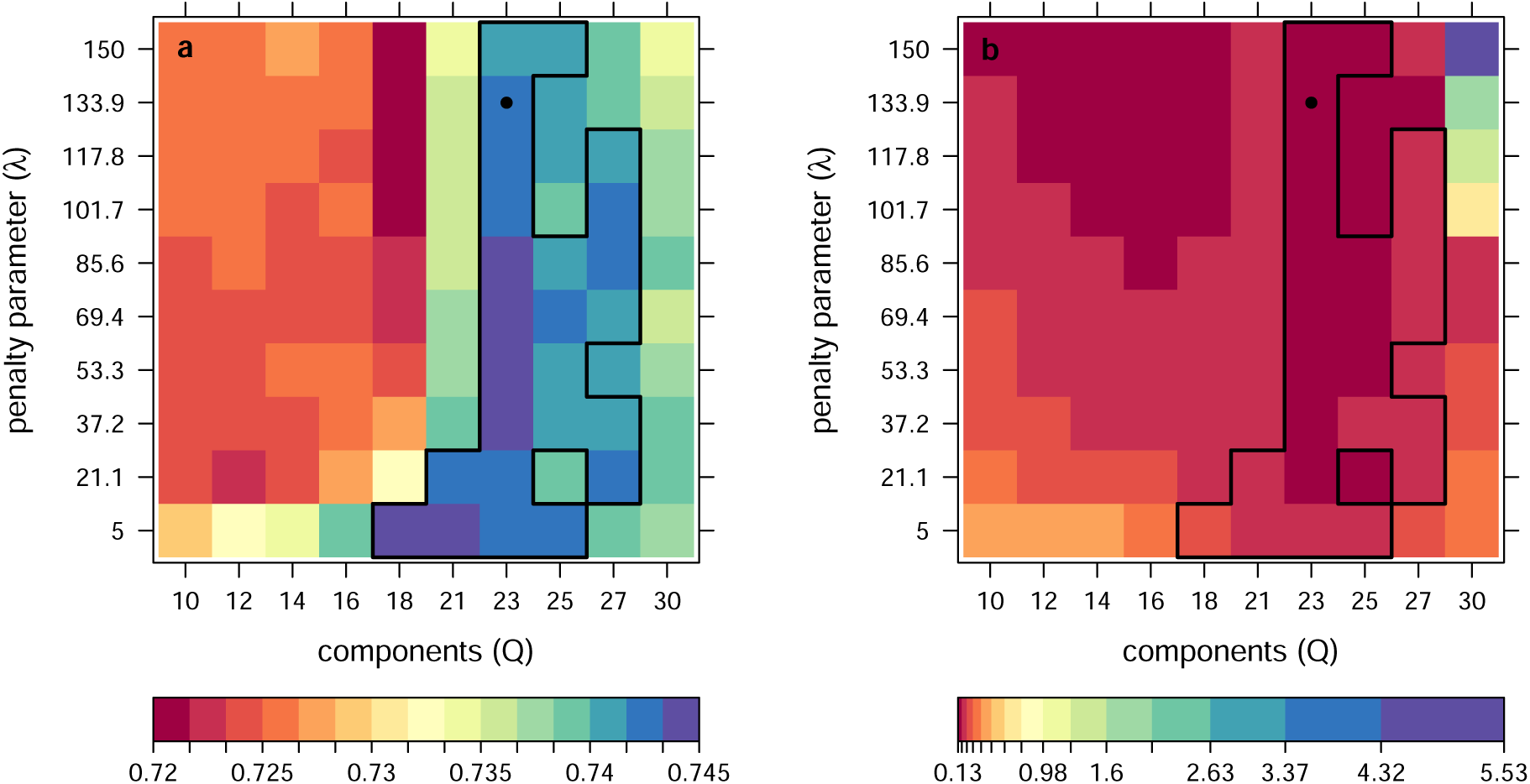
Maps showing (**a**) Area under the ROC curve (*AUC*) and (**b**) roughness of the coefficient function (*R*_*β*_), for `fspGLM` with different numbers of PLS components and penalty parameter, on the validating subset of the cross-validation dataset. The boxed area denotes the *n*_*a*_ models with acceptable performance, and the black circle denotes the smoothest model among those.

#### Diagnostics and results illustrated

We provide extensively annotated plots for quick assessment of model fit and prediction accuracy. These are designed to be as informative as possible so that potential issues with the training process can be easily identified. An example is shown in Figure 4 for a binary classification problem. Multinomial classification and regression problems have similar outputs.

**Figure 4:**
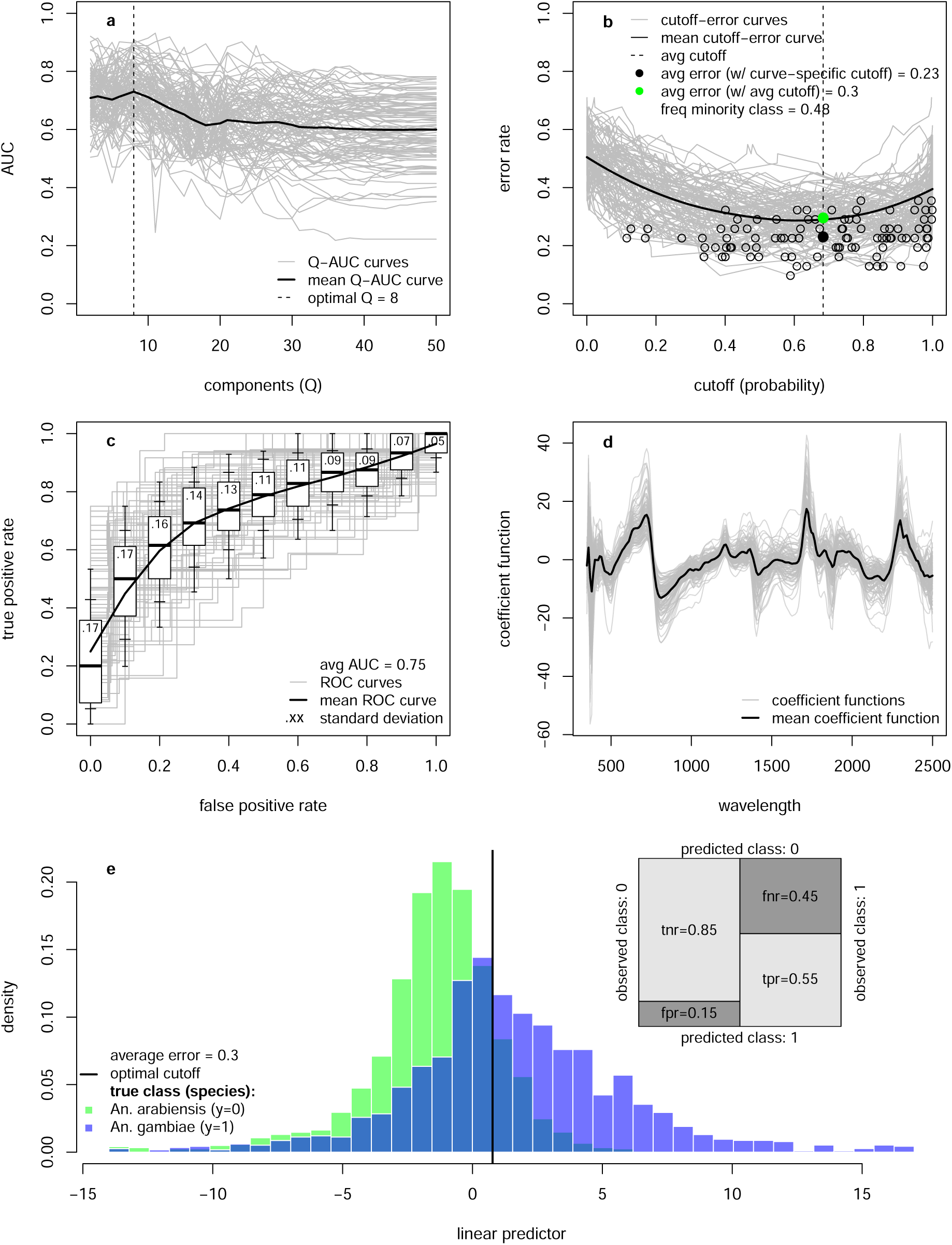
Diagnostic plots for a `fsGLM` with PLS components, showing (**a**) cross-validation for the number of components (*Q*); (**b**) cutoff-error plot displaying the choice of probability cutoff for classification; (**c**) ROC curves with variability shown in the boxplots (black line, box edges, inner and outer whiskers show 50th (median), 25th/75th, 15th/85th and 5th/95th percentiles, respectively); (**d**) coefficient function *β*(*t*); (**e**) histogram of the estimated linear predictor for the test observations, colour-coded by the true class. Results are averaged over 100 randomisations of the training/validating/testing subsets, shown individually in grey.

The *Q* cross-validation curves evaluated on the validating subset of the cross-validation dataset shows the performance achieved for a given number of PLS/PCA components (Figure 4a). When these curves are close to flat, it is useful to set a margin parameter 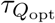 so that we choose not the *Q* corresponding to the largest AUC but one that gives an AUC within 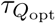 of the optimal. A small value such as 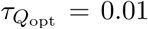 can often reduce the number of components substantially without considerable loss in performance, and furthermore can help prevent overfitting.

The classification cutoff–error curves depict the optimal probability cutoff that minimises the misclassification rate, giving equal weight to false positives and false negatives (Figure 4b). Models with optimal probability cutoff equal to 0 or 1 will always predict the same class, which can mean that a particular split of the dataset has come out highly imbalanced in terms of the responses classes and the model minimises the error by always predicting the majority frequency. Such models may deserve further investigation. The package also includes an option to enforce balanced dataset splits. An important quality requirement for any classifier using predictors (*C*_*p*_) is that it outperforms a naive classifier (*C*_*N*_) trained only on the frequency of the response variable (i.e., without predictors). The later will always predict the majority class, and so if our classifier outperforms this naive strategy we can be assured that the predictors contain valuable information. In other words, if the spectra is predictive of mosquito species, *C*_*p*_ must outperform *C*_*n*_. The color-coded dot in Figure 4b provides this information: green if *C*_*p*_ outperforms the *C*_*n*_; red otherwise. To fairly access performance we compute the misclassification rate using the average probability cutoff, although we also show the curve-specific cutoff for reference.

The ROC curves evaluated on the testing subset of the cross-validation dataset, together with dispersion measures and the AUC corresponding to the optimal classification cutoff as given in panel 4b give an overview of the performance of the model (Figure 4c).

The coefficient function *β*(*t*) can be used to identify the spectral regions of more importance for prediction, and is the key output of the model (Figure 4d).

A histogram of the estimated linear predictor for the test observations illustrates the models’ ability to separate the two classes (Figure 4e). The shaded area corresponds to misclassified observations, with false negatives to the left of the optimal cutoff line (*An. gambiae* incorrectly predicted to be *An. arabiensis*) and false positives to the right (*An. arabiensis* incorrectly predicted to be *An. gambiae*). The inset plot shows the confusion matrix with the breakdown of the classification results: true negative rate (tnr), false negative rate (fnr), true positive rate (tpr), false positive rate (fpr).

### 3.3 Identifying important predictors

Experiments to generate spectra from laboratory-reared mosquitoes remain relatively time-consuming, although necessary to train predictive models. With the view of simplifying and accelerating the process of gathering data, it is of interest to determine which variables under the experimenter’s control have an effect on spectra. The statistical framework presented, encapsulated in (8), can handle this type of hypothesis testing straightforwardly by testing the significance of the parameters ***γ*** associated with the non-functional predictors ***Z***.

To to illustrate this using the same dataset, we can determine whether the location of collection is a statistically significant. We use a binary variable `Location` encoding the location where samples were collected (Longo or Klesso). The average p-value for this variable in a `fsGLM` with balanced classes (*N* = 222) is *p* = 0.04, which provides some evidence that mosquitoes from the two collection locations have some differences which are not captured by the spectra. This result supports the use of penalised estimation and smoothing methods to prevent overfitting when doing cross-location prediction.

## 4 Discussion

NIRS has the potential to revolutionise entomological monitoring of mosquito-borne diseases though there is a need to refine the statistical methods used to translate spectral information into quantities of epidemiological interest. Spectra from mosquitoes with the same characteristics are also likely to vary from site to site reflecting the genetic heterogeneity in the mosquito population, local environmental factors and procedural differences between teams collecting and processing samples. If the NIRS is to become a widely used there is therefore a need to prevent statistical models converting spectra into mosquito characteristics to be generalizable and not overfitting to the local training dataset. Here we have identified a number of statistical techniques to support this process which should be adopted to increase the rigour of NIRS entomological monitoring. Spectra functional representation, spectra smoothing and penalisation for the coefficient function all improve the accuracy of NIRS models predicting mosquito species in the test dataset (independent mosquitoes collected from the same location) and more importantly on the alternative test dataset (mosquitoes collected 283km away). All the techniques provide a level of spectra smoothing, though the optimum use of these different methods (in combination or individually) will vary depending on the characteristics of the training and unknown dataset.

Smoother coefficient functions tend to give conservative estimates and therefore prevent overfitting to individual peaks which may result from measurement error generated by the machinery or represent a minor deviation specific to the local mosquito population. We would recommend all these methods are trialled with new and expanded datasets and the optimum model chosen using the methods outlined here.

The regularisation framework proposed here has several advantages over the standard methods used in the literature. The functional representation of spectra is more computationally efficient allowing models to be trained and fit quicker. Though this isn’t necessarily an issue with the dataset presented here (with single models being fit within a few minutes) this is likely to get more important as datasets grow and samples from multiple sites are used within the same model. Regularisation also provides a smoother coefficient functions which generalises better preventing overfitting to noisy spectra. This is particularly important when sample sizes are small and where instruments have high noise-to-signal ratio in some regions of the spectrum (for example at the ends their spectral ranges).

In these data both PCA and PLS as a method of dimension reduction tend to give similar results, but in some cases PLS requires fewer components than PCA to achieve a given accuracy as has been seen previously (de Jong, 1993a). In addition to the standard penalisation approach, the methods presented here also enable predictions to be made using either the smoothest or top 5 smoothest models selected from the best performing models. Here they were selected for by choosing the models with either the smoothest or the 5 smoothest coefficient functions which were drawn from the top 25 most accurate models as evaluated on the validating subset of the cross-validation dataset (from mosquitoes within the same village). In these data this did not substantially improve the accuracy when predicted the species of the second village. This confirms the robustness of the strategy employed here of selecting the most appropriate smoothing method in order to obtain good generalizability. Further work with larger more diverse datasets are needed to understand the benefit of selecting the smoothest over the best fitting models. Here we defined an acceptably accurate model as being within 1 percentage point of the most accurate model, although this parameter will need to be refined according to the question under investigation which will determine the trade-off between accuracy and generalisability. Similarly, the ensemble method proposed here which selected the 5 smoothest models from the top 25 most accurate did not perform better than the single best fit model. The added benefit of this ensemble approach needs to be investigated further using larger datasets collected from more diverse geographical locations as it may be expected to perform better in these scenarios.

Our results indicate that the accuracy of NIRS ability to determine the sibling species of mosquito within the An. gambiae complex is lower than previous estimates. This work was intended to showcase the different statistical methods and not evaluate the technique and there are a number of reasons why the moderate accuracy should not be overly interpreted. Firstly, the sample size used in this study is very small with only 126 samples available. This means that only 63 samples were used to train each model (a different set of 63 samples for each of the 100 models), which is a very low number given the diversity of spectra. Future work may have sample sizes an order or two of magnitude larger if the technique is adopted further. Secondly, there were a mixture of F0 and F1 mosquitoes used in this analysis. Spectra collected from mosquitoes which were caught in different ways may vary, though the number of mosquitoes available for this analysis was insufficient to test this here. Lastly, other characteristics of interest which might influence spectra were not collected or included in the model. For example the level of insecticide resistance may vary substantially within the same species within the same population and has been shown to influence specta (). It is worth noting however that high accuracy isn’t necessarily a prerequisite for NIRS to be a useful tool (Lambert et al., 2018). NIRS could also be used as a pre-scanning tool, for instance to determine if mosquitoes are infected before parasite genetic sequencing. In that case, we are interested in maximising the ratio of truly infected to truly uninfected TP/(TP +FP) in order to minimise the cost per mosquito sequenced. This can be done with ROCR package by choosing the ppv (positive predictive value) criterion in the function performance(). Additionally, there can be class imbalance which leads to imbalanced misclassification rates. Tuning the importance of false positive rates to false negative rates can help giving balanced misclassification rates for the different classes according to the question under investigation.

## Acknowledgements

The work was supported by UK Medical Research Council (MRC) Project Grant (MR/P01111X/1). We acknowledge joint Centre funding from the UK Medical Research Council and Department for International Development (MR/R015600/1).

## Appendix

### A Dimension reduction

#### Functional principal component analysis (fPCA)

Principal component analysis was originally used to overcome multicollinearity in the linear model (Jolliffe, 2002; Massy, 1965) and subsequently extended to functional data (Cai and Hall, 2006; Cardot et al., 1999; Hall and Hosseini-Nasab, 2006; Müller and Stadtmüller, 2005; Shang, 2014; Wang et al., 2016). The objective of fPCA is to compute directions that maximise the variance of the functional data *X*(*t*) when projected onto these directions. Assuming E[*X*(*t*)] = 0, ∀*t*∈ 𝒯 for notational simplicity, we can formalise fPCA as solving the following problem:

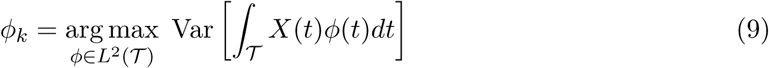

subject to ‖*ϕ*‖ = 1 (normalisation) and ∫_𝒯_ *ϕ*_*l*_(*t*)*ϕ*(*t*) = 0, ∀*l* < *k* (orthogonality). Here, *ϕ*_*k*_ is the *k*th orthogonal fPCA direction (*loading*) associated with the covariance function *K*(*s, t*) = Cov[*X*(*s*), *X*(*t*)]; and *v*_*ik*_ = ∫_𝒯_ *X*_*i*_(*t*)*ϕ*_*k*_(*t*)*dt* is the *k*th fPCA component (*score*), that is, the projection of *X*_*i*_ onto *ϕ*_*k*_. By construction, *ϕ*_1_ gives the direction of highest variation; *ϕ*_2_ the direction of next highest variation that is orthogonal (uncorrelated) to *ϕ*_1_; and so on.

Even in high dimensional data, often a small number of these components is sufficient to capture most of the variation in *X*. This feature selection procedure can therefore accommodate dimension reduction with minimal information loss, the trade-off being regulated by the tuning parameter *Q*, which can be chosen by cross-validation.

##### Computation

Following (5), we derive the fPCA components not from ***X*** but from ***XB***. The fPCA components are estimated by singular value decomposition, ***XB*** = ***U* Σ*V*** ^*T*^. This produces a matrix ***V*** whose columns [***v***_1_, ***v***_2_, …,] are the fPCA components or eigenvectors of the covariance matrix of ***XB***, approximating the eigenfunctions [*ϕ*_1_, *ϕ*_2_, …]. The dimension reduction projection matrix in (8) is then ***D*** = ***V***_*Q*_, where ***V***_*Q*_ denotes the matrix whose columns are the first *Q* columns of ***V***.

#### Functional partial least squares (fPLS)

Partial least squares was also originally used to solve multicollinearity among predictors in the context of linear models, and has been widely used in chemometrics (Geladi and Kowlaski, 1986; Wold et al., 1984, 2001) and extended to functional data (Aguilera et al., 2010; Delaigle and Hall, 2012; Preda and Saporta, 2005). The objective of fPLS is to identify directions which maximise the covariance between the response *y* and the functional data *X*(*t*) when projected onto those directions:

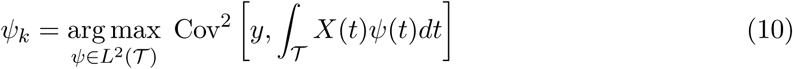

subject to ‖*ψ*‖ = 1 (normalisation) and ∫_𝒯_ ∫_𝒯_ *ψ*_*l*_(*s*)Σ(*s, t*)*ψ*(*t*)*ds dt* = 0, ∀*l* < *k* (covariance-orthogonality), where Σ(*s, t*) denotes the covariance function of *X*. Here, *ψ*_*k*_ is the *k*th covariance-orthogonal fPLS direction; and *r*_*ik*_ = ∫_𝒯_ *X*_*i*_(*t*)*ψ*_*k*_(*t*)*dt* is the *k*th fPLS component, that is, the projection of *X*_*i*_ onto *ψ*_*k*_.

The interpretation of the sequential optimisation problem is similar to the case of fPCA, except that fPLS maximises the covariance between response and predictor instead of the predictor variance. This addresses an important concern, namely that in fPCA the response is not considered and, therefore, there are no guarantees that the components explaining the most variation in the functional predictor are also the best at explaining the relation between the predictor and the response—which is the ultimate goal of the analysis—although the two tend to be related (de Jong, 1993a; Mevik and Wehrens, 2015).

##### Computation

As before, we derive the fPLS components from ***XB***. Several algorithms have been proposed, for example NIPALS (Wold et al., 1984) or SIMPLS (de Jong, 1993b). These produce a matrix ***R*** whose columns [***r***_1_, ***r***_2_, …,] are the fPLS components approximating [*ψ*_1_, *ψ*_2_, …]. The dimension reduction projection matrix in (8) is then ***D*** = ***R***_*Q*_, where ***R***_*Q*_ denotes the matrix whose columns are the first *Q* columns of ***R***.

